# A chromosome-level genome assembly of *Zasmidium syzygii* isolated from banana leaves

**DOI:** 10.1101/2023.08.24.554682

**Authors:** A.C. van Westerhoven, R. Mehrabi, R. Talebi, M.B.F. Steentjes, B. Corcolon, Pablo A. Chong, G.H.J. Kema, M. F. Seidl

## Abstract

Accurate taxonomic classification of samples from infected host material is essential for disease diagnostics and genome analyses. Despite the importance, diagnosis of fungal pathogens causing banana leaf diseases remains challenging. Foliar diseases of bananas are mainly caused by three *Pseudocercospora* species, of which the most predominant causal agent is *P. fijiensis*. Here, we sequenced and assembled four fungal isolates obtained from necrotic banana leaves in Bohol (Philippines) and obtained a high-quality genome assembly for one of these isolates. The samples were initially identified as *P. fijiensis* using PCR diagnostics, however, the assembly size was consistently 30 Mb smaller than expected. Based on the ITS sequences, we identified the samples as *Zasmidium syzygii* (98.7% identity). The high-quality *Zasmidium syzygii* assembly is 42.5 Mb in size, comprising 16 contigs, of which 11 are complete. The genome contains 98.6% of the expected single-copy BUSCO genes and contains 14,789 genes and 10.3% repeats. The three short-read assemblies are less continuous but have similar genome sizes (40.4 – 42.4 Mb) and contain between 96.5% and 98.4% BUSCO genes. All four isolates have identical ITS sequences and are distinct from *Zasmidium* isolates that were previously sampled from banana leaves. We thus report the first continuous genome assembly of a member of the *Zasmidium* genus, forming an essential resource for further analysis to enhance our understanding of the diversity of pathogenic fungal isolates as well as fungal diversity.

## Introduction

Genome sequencing is important for the diagnostics and monitoring of diseases and is an important step to understand the biology of pathogens and diseases. To enable sample identification and downstream genome analyses, accurate taxonomic classification and prevention of contamination in genome assemblies are essential (Francois et al., 2020; Lu & Salzberg, 2018; Rachtman et al., 2020). However, publicly available genome assemblies are occasionally reported to contain contaminants, which can lead to incorrect species classification (Cornet & Baurain, 2022; Kusch et al., 2023; Steinegger & Salzberg, 2020). Obtaining a clean genome assembly is especially challenging for pathogens that live in close association with their host; samples from these pathogens are often contaminated with host material or other organisms that proliferate in proximity to the host such as endophytic fungi (Kusch et al., 2023; Zaccaron & Stergiopoulos, 2021).

Banana is an important food crop providing food security in tropical and subtropical regions worldwide. Foliar blights are a major constraint to banana production and are mainly caused by a complex of three *Pseudocercospora* species with *P. fijiensis* as a major constituent that causes black leaf streak disease or black Sigatoka. Control of this disease is responsible for up to 25% of the total costs of banana production (Drenth & Kema, 2021). The two other species also cause foliar blights, but are currently less prevalent, *P. musae* causes yellow Sigatoka and *P. eumusae* causes eumusae leaf spot (Chang et al., 2016). However, besides these three *Pseudocercospora* species, other fungal species can appear in association with symptomatic banana foliage. For example, a recent study that analysed fungal isolates associated with banana foliar diseases revealed the presence of over 30 other fungal species, primarily belonging to the *Mycosphaerellacea* family (Crous et al., 2021). Interestingly, before this study revealed the identity of these isolates, most of these fungal species were considered to belong to the *Pseudocercospora* genus, highlighting that accurate diagnosis of the causal agents of leaf symptoms remains challenging. Yet, accurate identification and classification of the obtained isolates is important, particularly for developing disease diagnostics and enhancing the effectiveness of disease management strategies (Kusch et al., 2023; Lu & Salzberg, 2018).

Here we sampled four fungal isolates from banana foliage showing blight symptoms in Bohol, Philippines, and identified the fungal isolates by disease symptoms, morphology, and diagnostic PCR assays (Arzanlou et al., 2008). Interestingly, initial observations classified the isolates as *P. fijiensis*, but further genome analyses revealed that these isolates rather represent *Zasmidium syzygii*. Until now, only three fragmented genome assemblies of *Zasmidium* species are publicly available (Haridas et al., 2020; Xu et al., 2017). Here, we assembled the first chromosome-level genome assembly of a representative of the *Zasmidium* genus. The availability of this genome assembly will help to improve molecular diagnostics for pathogenic *Pseudocercospora* spp. on banana foliage. Moreover, the genome adds to the diversity of available fungal genomes, providing a resource for future genomic studies.

## Methods & Materials

### Fungal isolation and sequencing

Banana leaves with symptoms of foliar disease were collected from a field in Bohol, Philippines. Leaf samples were cut into 1 cm^2^ segments and used to discharge ascospores as described previously (Chong et al., 2019). Four single ascospore isolates (P121, P122, P123, and P124) were collected and grown on potato dextrose agar (PDA) plates supplemented with streptomycin (100 μg/ml) for three weeks at 25 °C. To obtain sufficient fungal biomass for DNA isolation, a piece of mycelium (2 cm^2^) was blended for 20 sec. at 6,000 rpm in an Ultra Turrax Tube Drive homogenizer (IKA, Staufen, Germany) in 15 ml water using a sterile DT-20 tube (IKA, Staufen, Germany). The fragmented mycelium was transferred into a flask containing 100 ml of PDB amended with streptomycin (100 µg/ml) and kept at 25 °C on a rotary shaker with 150 rpm for about two weeks. The fungal mycelium was filtered through miracloth and subsequently washed with sterile water. Fungal mycelium was freeze-dried overnight and used for high-molecular-weight (HMW) DNA isolation based on the CTAB method (Murray & Thompson, 1980). After adding isopropanol, HMW DNA was collected from the extraction buffer using a sterile needle. AMPure XP purification kit (Becman Cluter Life Sciences, USA) was used to clean up the DNA. DNA quality and quantity were checked by gel electrophoresis, Nanodrop micro-volume spectrophotometers, and Qubit Fluorometric Quantitation (Thermo Fisher Scientific, USA). The HMW DNA of isolate P124 was sequenced using PromethION Oxford Nanopore (ONT) sequencing technologies. Additionally, all isolates were sequenced using the Illumina HiSeq platform, both sequencing platforms were located at Keygene B.V. (Wageningen, the Netherlands).

### Fungal species diagnosis by PCR

To identify the isolated fungal species, the DNA from each isolate was extracted and subjected to *P. fijiensis*-specific diagnostic PCR according to a previously established protocol (Arzanlou et al., 2008). The list of primers used in PCRs is shown in Table S1. Actin (820 bp) was used as a positive control to ensure successful amplification and to assess the quality of DNA.

### Pathogenicity assay

The pathogenicity of isolate P124 was tested on two banana genotypes; Cavendish cv. Grand Naine (AAA) and a diploid (AA) genotype ‘Pisang Berlin’. The inoculum for isolate P124 as well as for the *P. fijiensis* reference isolate P78 were prepared similarly. A piece of 1 cm^2^ mycelium from three-week-old colonies grown on PDA was collected in an Eppendorf tube containing 3-4 metal beads (3 mm diameter) and was blended for 20 seconds at 3,000 rpm in a Bead Beater Homogenizer. About one ml of sterile water was added to each tube and fragmented mycelium was spread on PDA plates amended with 100 µg/ml streptomycin. The plates were kept at 25°C for 3-4 weeks and then a piece of mycelium (10 cm^2^) was blended for 40 sec. at 6,000 rpm in an Ultra Turrax Tube Drive homogenizer (IKA, Staufen, Germany) in 15 ml of distilled water using a sterile DT-20 tube (IKA, Staufen, Germany). The suspension was passed through miracloth to remove non-fragmented mycelium. The collected mycelial fragments were further diluted and adjusted to 5×10^5^ fragments ml^-1^ and supplemented with 0.15% Tween 20. This suspension was used to inoculate two-month-old banana plants on both sides of the leaves. Each treatment was repeated tree times, as a control, water was used for mock inoculation. Inoculated plants were kept for 48 h at 90% relative humidity at 25⁰C in the dark in a growth cabinet and subsequently for eight weeks in a greenhouse with >85% RH, and with a day length of 12 hours.

### De novo genome assembly

To construct a complete and continuous genome assembly, we assembled the ONT long-read sequences of isolate P124. The raw Fast5 files were basecalled with Guppy (v. 5.1.13) and filtered for a minimum quality score of 10. The filtered reads were assembled *de novo* using Canu (v. 2.2, Koren et al., 2017) with the parameters ‘-OvlMerThreshold=300’ and ‘-corMaxEvidenceErate=0.15’ to improve the assembly of repetitive regions. The initial genome assembly was polished with Racon (v. 1.5.0 Vaser et al., 2017) using the raw ONT long reads. After polishing, missing telomeres were reconstructed, when possible, using teloclip (v. 0.0.3, https://github.com/Adamtaranto/teloclip) and a final polishing step was performed based on shrot-read Illumina data using Pilon (v. 1.24, Walker et al., 2014). The presence of telomeres (TTAGGG) and coverage of the chromosomes in the final polished assembly were determined using tapestry (v. 1.0.0, Davey et al., 2020). To estimate the completeness of the genome assembly, the presence of conserved single-copy genes was determined using BUSCO with the Capnodiales database (Simão et al., 2015). The assembly continuity and potential contaminations were accessed using BlobTools (v. 4.1.5, Challis et al., 2020). To estimate the size of the assembled genome, all Illumina sequences were analyzed using GenomeScope (v. 2.0, Vurture et al., 2017) with the 21-mer profile determined by JellyFish (v 2.3.0, Marçais & Kingsford, 2011). Isolate P121 showed high k-mer coverage outliers, therefore the maximal k-mer coverage was set to 350x, similar to the maximum k-mer coverage observed in the other genome. Repeats were predicted using RepeatModeler (v. 2.0.3, Flynn et al., 2020) and annotated using RepeatMasker (v. 4.1.2, Smit et al., 2015). Finally, protein-coding genes were annotated using Funannotate (v. 1.8.9, Palmer & Stajich, 2019) with the BUSCO lineage pezizomycotina utilizing predicted *Zasmidium cellare* proteins as evidence (GCF_010093935.1, Haridas et al., 2020). Additionally, we assembled the Illumina sequenced isolates P121-P123 with Spades (v. 3.13, Bankevich et al., 2012). To assess the quality and completeness of these assemblies, we ran QUAST (v. 5.2, Gurevich et al., 2013) and BUSCO (v. 5.3.2, Simão et al., 2015) with the Capnodiales odb10 database.

### Identification of internal transcribed spaces sequence

The internal transcribed spacer (ITS) sequences from the four assemblies (P121-P124) were retrieved using ITSx (v. 1.1.3, Bengtsson-Palme et al., 2013) and blastn was used to detect similar ITS sequences in the blast database (accessed 19^th^ November 2022). Additionally, a maximum-likelihood tree was constructed to compare the four ITS sequences to 227 publicly available ITS sequences obtained from NCBI (accessed 13^th^ January 2023). Sequences were aligned using mafft (v.7. 453, Katoh et al., 2002) and a maximum-likelihood phylogenetic tree was constructed using RAxML, with 500 bootstrap replicates (-m GTRCAT -p 1234 -b 100 -N 500) (v. 8.2.12, Stamatakis, 2014). The resulting phylogeny was visualized using iTOL (Letunic & Bork, 2021).

### Genetic diversity

To assess the genetic diversity among isolates, short-read sequencing data of P121-P123 were aligned to the P124 long-read genome assembly using BWA-mem (v. 0.7.17, Li, 2013). GATK4 was used to call variants (Van der Auwera et al., 2013), and these variants were subsequently filtered using GATK Variant Filtration based on the GATK best practices (Van der Auwera et al., 2013). The filtering process involved excluding variants from reads with low mapping quality, variants predominantly located at the edge of reads, and variants exhibiting a bias towards reverse/forward strands.

## Results and Discussion

### Fungal isolates obtained from symptomatic banana foliage in Bohol

Foliar blights of banana is mainly caused by a complex of *Pseudocercospora* species (Chang et al., 2016; Drenth & Kema, 2021). However, additional fungal species predominantly belonging to the *Mycosphaerella* genus have also been associated with symptomatic banana leaves (Crous et al., 2021). To further analyze the fungal pathogens that cause disease on banana leaves, four samples were obtained from banana plants with necrotic lesions in Bohol, Philippines (Chong et al., 2021). For all four isolates (P121 – P124) we were able to amplify a PCR product using *P. fijiensis*-specific PCR primers (Arzanlou et al., 2008), indicating that the isolates can be identified as *P. fijiensis* (**Figure 1a**). To corroborate the identity of the isolates, we tested the pathogenicity of one representative (isolate P124) on banana cultivars Cavendish (cv. Grand Naine AAA) and Pisang Berlin (AA) and compared the pathogenicity to the reference *P. fijiensis* isolate P78 originating from Tanzania. No leave spot symptoms were observed on Cavendish upon inoculation with P124, in contrast to the necrotic lesions caused by P78 (**Figure 1b**). However, both isolates caused necrotic lesions on Pisang Berlin four weeks after inoculation (**Figure 1c**). The necrotic lesions caused by isolate P124 were less severe than the disease symptoms caused by *P. fijiensis* isolate P78. Although we observe a difference in pathogenicity between the two isolates differ in pathogenicity, both isolates but cause necrosis on banana foliage.

**Figure 1.**
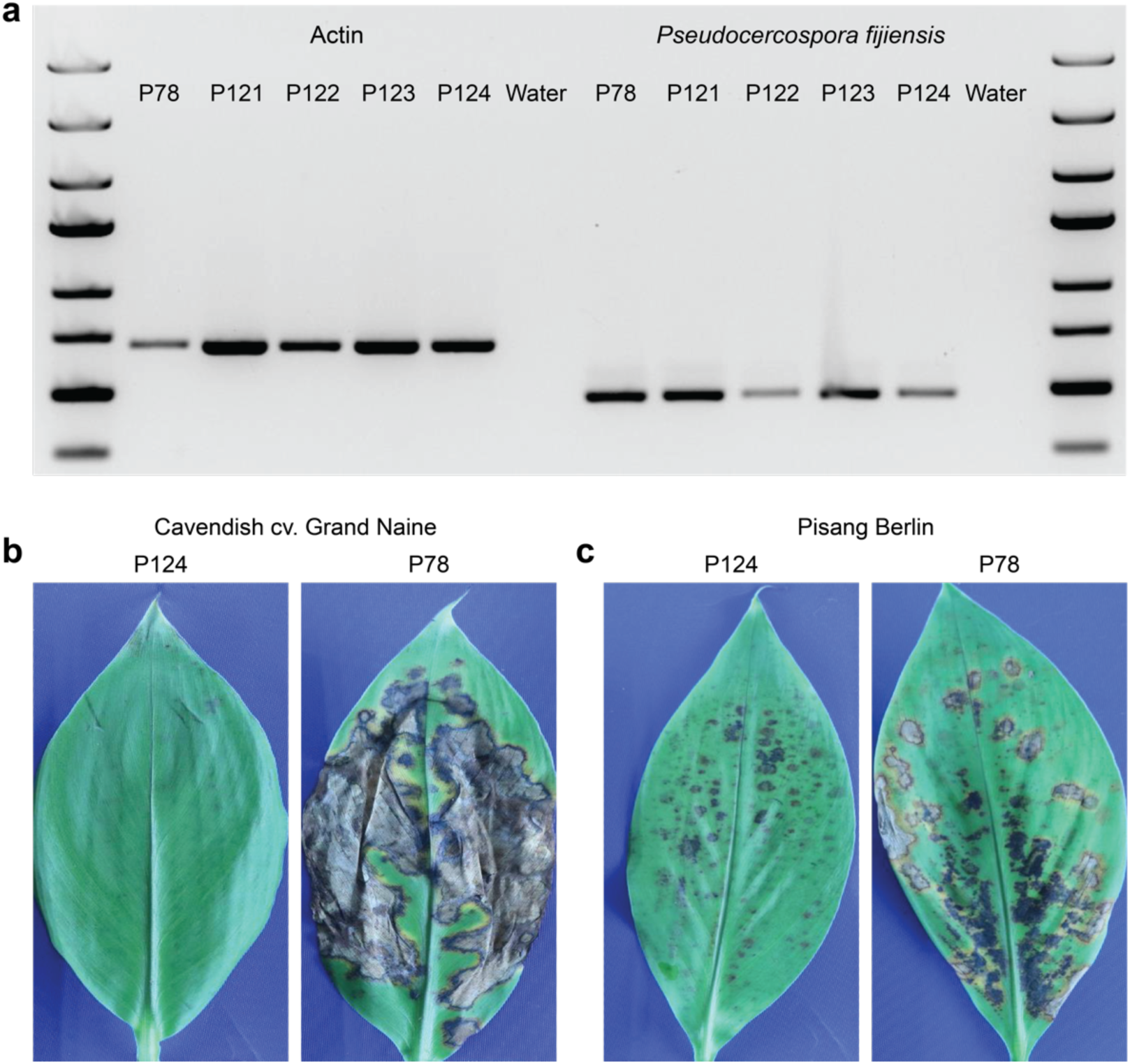
PCR diagnostics and pathogenicity assay of isolate P124 in comparison to the *Pseudocercospora fijiensis* reference strain P78. a) All isolates show amplification with actin primers (left panel). *Pseudocercospora*-specific primers amplified PCR products from *P. fijiensis* strain P78 (positive control) as well as from isolates P121, P122, P123, and P124 (right panel), suggesting that the isolates can be identified as *P. fijiensis*. b) Isolate P124 does not cause necrotic lesions on Cavendish banana, in contrast to necrotic symptoms caused by P78. c) Both P124 and P78 cause necrotic leaf symptoms on Pisang Berlin. Disease symptoms were scored eight weeks after inoculation.

### Chromosome-level genome assembly of P124

To generate a high-quality genome assembly for the isolates from Bohol, we randomly selected strain P124 for sequencing using Oxford Nanopore Technology. This yielded 11.9 Gb of reads with an average size of ~9.1 kb and a read N50 of 12 kb, corresponding to a ~150x genome coverage based on the estimated genome size of 74 Mb (Arango et al., 2016). The *de novo* genome assembly resulted in 16 nuclear contigs and one contig representing the mitochondrial genome. The assembly has an N50 of 3.4 Mb and a total genome size of 42.5 Mb. Eleven of the 16 contigs had telomeric sequences (TTAGGG) at both ends, and thus the assembly is highly contiguous and mostly represents complete chromosomes (Figure 2). To assess genome completeness, we queried the genome assembly for the presence of single-copy BUSCO genes and identified 98.6% of the single-copy BUSCO genes that are expected in the fungal order Capnodiales, indicating that the genome assembly of P124 covers the conserved gene space (Figure 2a). We predicted a total of 14,789 protein-coding genes in the genome and identified that 10.3% of the genome consists of repetitive elements (Figure 2b). *De novo* genome assembly of isolates P121 – P123, sequenced with short-read sequencing technology only, resulted in genome assemblies with similar sizes (40.4 – 42.5 Mb). Although the assemblies of P121 - P123 are more fragmented compared to the assembly of the nanopore-sequenced isolate P124, they approximately contain an equally high number of expected BUSCO genes (Table 1).

**Figure 2.**
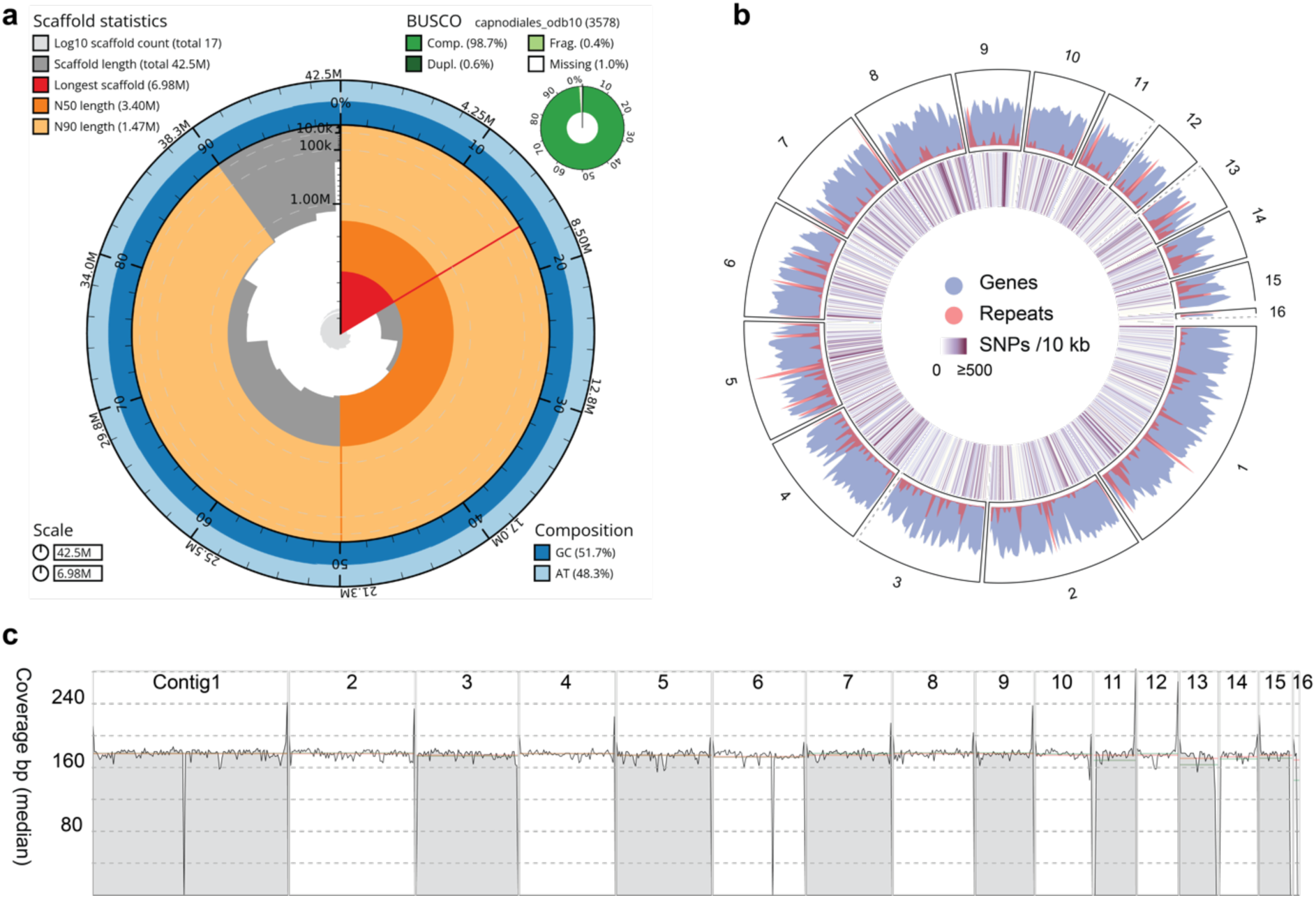
Chromosome-level genome assembly of *Zasmidium syzygii* isolate P124. **a**) Genome assembly statistics for the *de novo* assembly of *Z. syzygii* isolate P124 based on Oxford Nanopore Technology (ONT). The genome assembly has 17 contigs (16 nuclear contigs of which at least 11 are complete chromosomes, and the mitochondrial genome) with a total genome size of 42.5 Mb and contains 98.7% complete single-copy BUSCO genes. **b**) A circular representation of the contigs in P124. Dashed lines indicate missing telomeres for chromosome 3, 11, 13, 15, and 16. Genes (14,786) and repeats (10.3%) are distributed evenly over the chromosomes. Single nucleotide polymorphisms are found over all contigs, with two SNP dense regions on contig 8 and 9. **c**) Short-read coverage of P124 mapped to the assembled contigs of P124 shows minimal regions with exceptionally high or low coverage, suggesting that the assembly does not contain large repetitive regions that may have been collapsed during the assembly process. Mean coverage (green line) and Median coverage (red line) are indicated in the figure.

**Table 1.** The genome assembly statistics of four sequenced isolates obtained from symptomatic banana foliage in Bohol, Philippines.

|  | **P124\*** | **P121\*\*** | **P122\*\*** | **P123\*\*** |
| --- | --- | --- | --- | --- |
| Genome size (Mb) based on k-mer estimation (k=21) | 40.6 | 39.8 | 39.9 | 41.9 |
| Genome assembly size (Mb) | 42.5 | 40.4 | 40.6 | 42.2 |
| Number of contigs | 16 | 1,225 | 1,563 | 796 |
| Largest contig (Mb) | 6.97 | 0.44 | 0.31 | 0.84 |
| GC% | 51.70 | 52.55 | 52.37 | 52.01 |
| N50 (Mb) | 3.40 | 0.98 | 0.60 | 1.9 |
| N75 (Mb) | 2.08 | 0.49 | 0.30 | 1.05 |
| L50 | 2 | 134 | 201 | 67 |
| BUSCO completeness ($n = 3,578$) | 98.6% | 97.8% | 96.5% | 98.4% |
| BUSCO fragmented genes | 0.5% | 0.9% | 1.5% | 0.5% |
| BUSCO missing genes | 1.0% | 1.3% | 2.0% | 1.1% |

\*P124 is assembled using long reads from Oxford Nanopore Technology (ONT).
\*\*P121-P123 are assembled using Illumina short reads.

|  | **P124\*** | **P121\*\*** | **P122\*\*** | **P123\*\*** |
| --- | --- | --- | --- | --- |
| Genome size (Mb) based on k-mer estimation (k=21) | 40.6 | 39.8 | 39.9 | 41.9 |
| Genome assembly size (Mb) | 42.5 | 40.4 | 40.6 | 42.2 |
| Number of contigs | 16 | 1,225 | 1,563 | 796 |
| Largest contig (Mb) | 6.97 | 0.44 | 0.31 | 0.84 |
| GC% | 51.70 | 52.55 | 52.37 | 52.01 |
| N50 (Mb) | 3.40 | 0.98 | 0.60 | 1.9 |
| N75 (Mb) | 2.08 | 0.49 | 0.30 | 1.05 |
| L50 | 2 | 134 | 201 | 67 |
| BUSCO completeness ($n = 3,578$) | 98.6% | 97.8% | 96.5% | 98.4% |
| BUSCO fragmented genes | 0.5% | 0.9% | 1.5% | 0.5% |
| BUSCO missing genes | 1.0% | 1.3% | 2.0% | 1.1% |

\*P124 is assembled using long reads from Oxford Nanopore Technology (ONT).
\*\*P121-P123 are assembled using Illumina short reads.

The *P. fijiensis* reference genome is 73.6 Mb (Arango et al., 2016), and remarkably, the genome assemblies of the here sequenced fungal isolates are approximately 30 Mb smaller than expected. To verify the assembly size, we estimated genome sizes using k-mer profiles, which resulted in an estimated genome size ranging from 38.9 to 41.9 Mb, similar to the size of the *de novo* assembled genomes (Table 1). Furthermore, we mapped the sequencing reads of isolates P121 – P124 to the chromosome-level genome assembly of P124, mapping on average 95% of the reads with limited genetic diversity between isolates (on average 15 SNPs per kb; Figure 2b). An increased read mapping coverage is observed in most telomeric regions (Figure 2c), while the coverage drops in contig 1, contig 6, and contig 16, suggesting that these regions possibly contain assembly artefacts. Contig 16 is only 0.2 Mb in size and only contains a telomeric repeats at one end. The low coverage region together with read mappings that support a possible link to either contig 3 or 14, which also lack one of the telomeric repeats, suggests that this contig might be associated to one of these two contigs. Apart from these regions, mapping of the short reads of P124 to the P124 chromosome-level genome assembly revealed an overall constant read mapping coverage over the chromosomes, indicating that the P124 genome assembly does not contain extensively collapsed repetitive regions that could account for the smaller genome size compared to *P. fijiensis*. This validates that, although the genome assembly of strain P124 is smaller than expected, it is highly complete and continuous, suggesting that the length variation is likely not due to assembly errors. To capture possible contaminations in the genome assemblies, we queried all contigs in the BLAST database to determine their identity. All contigs showed a high similarity to members of the *Mycosphaerellacea* family; most contigs were similar to *Zasmidium* species (13 contigs, 35 Mb) and only one contig (3.3 Mb) showed similarity to *Pseudocercospora* species (**Figure S1**). Whole-genome alignments of our assembly to the *P. fijiensis* reference genome assembly revealed that none of the contigs display significant similarity (**Figure S2**). Therefore, we considered that the difference in genome size between isolates P121 – P124 and the *P. fijiensis* reference isolate P78 is likely caused by the isolation of a different fungal species associated with necrotic symptoms on banana foliage (Crous et al., 2021), most likely by a member of genus *Zasmidium*.

### The assembled genome sequence reveals that isolate P124 belongs to the fungal genus *Zasmidium*

To determine the identity of isolate P124, we retrieved the ITS sequence from the genome assembly and searched for related species in the non-redundant BLAST database on NCBI using the P124 ITS sequence as a query. We obtained a highly similar match (98.7% nucleotide identity) to *Zasmidium syzygii* (NR_111826.1), supporting that the assembled isolate belongs to the *Zasmidium* genus. In line with this finding, previous studies have found *Zasmidium* isolates on symptomatic banana leaves (Arzanlou et al., 2008; Crous et al., 2021), and other *Zasmidium* species are reported to cause leaf spot diseases on other plant species such as citrus (Aguilera-Cogley et al., 2017; An et al., 2021; Han et al., 2015). The *Zasmidium* genome assembly we report here is less fragmented and 4 Mb larger than the previous genome assembly of a *Zasmidium* species; *Z. cellare*. The species, known as the ‘wine cellar fungus’ because it thrives in walls and ceilings of wine cellars (Tribe et al., 2006), was sequenced and its genome was assembled into >267 scaffolds with a genome size of ~38 Mb (Haridas et al., 2020), illustrating that our genome offers a more continuous genome representation of a member of the *Zasmidium* genus.

To validate the identity and to determine the diversity of the *Zasmidium* isolates, we compared the ITS sequences of isolates P121-P124 with 43 other *Zasmidium* ITS sequences from NCBI (20 December 2022) as well as ITS sequences of 120 *Mycosphaerella* strains obtained from banana leaves (Crous et al., 2021). A maximum-likelihood phylogeny based on the aligned ITS sequences confirmed that isolates P121 – P124 belong to the *Zasmidium* genus and shows that these four isolates encode identical ITS sequences (Figure 3). Notably, the set of ITS sequences also contains sequences from five other *Zasmidium* isolates that had been previously sampled from banana leaves from different geographic locations (Martinique, Tonga, and Gabon) (Crous et al., 2021). Interestingly, these do not cluster with isolates P121-P124, suggesting that pathogenicity towards banana is a polyphyletic trait within the *Zasmidium* genus (Figure 3). Although *Zasmidium* species have been linked to foliar blights in various hosts (Aguilera-Cogley et al., 2017; Han et al., 2015; Osorio et al., 2021), the pathogenicity and global spread of *Zasmidium* species has not been studied in depth. Based on our data, we conclude that Z. *syzygii* occurs on banana leaves with necrotic symptoms and can be the cause of mild necrotic lesions on the foliage. However, the abundance and role of Z. *syzygii* as a banana pathogen remains unknown, which requires further research to understand its prevalence, significance, and potential impact on banana cultivation.

**Figure 3.**
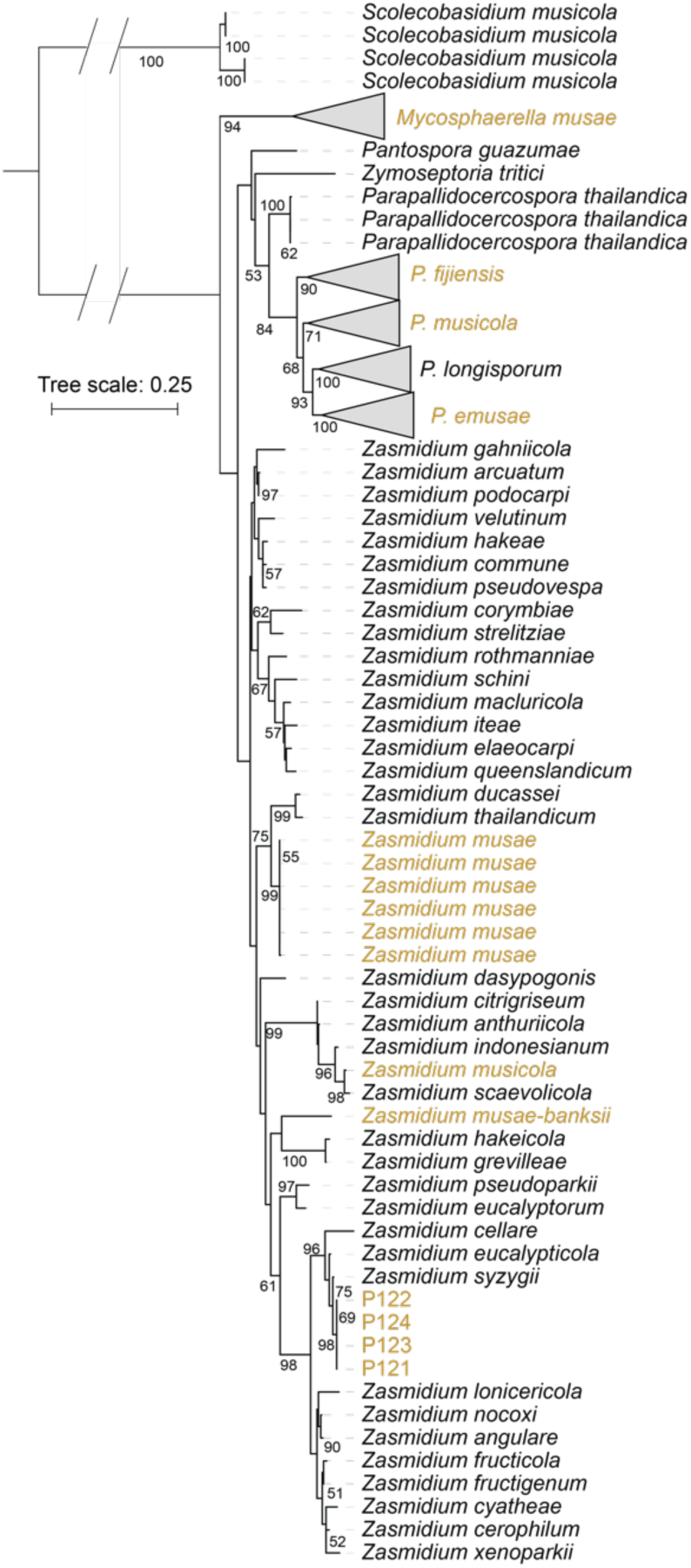
The four isolates sampled from Bohol (P121-P124) belong to the genus *Zasmidium.* A maximum-likelihood phylogenetic tree was constructed using the internally transcribed spacer (ITS) sequences of 43 *Zasmidium* species from NCBI and 120 *Mycosphaerellacea* isolates from banana leaves (Crous et al., 2021). The ITS sequence extracted from the chromosome-level genome assembly (P124) as well as from the other isolates (P121-123) associated with an isolate classified as *Z. syzgii* and are not related to *Zasmidium* strains previously isolated from banana leaves. *Zasmidium* isolates associated with infected banana leaves (yellow labels) are genetically diverse and are distributed across various branches of the phylogenetic tree.

Accurate diagnostics of pathogens is essential to detect the emergence and trace the dispersal of diseases, which is pivotal for effective disease management. However, our data reveal that *P. fijiensis* and *Z. syzygii* are indistinguishable with the current PCR diagnostic (Arzanlou et al., 2008). To compare the similarity of the PCR primers between *P. fijiensis* and *Z. syzygii*, we *in silico* detected the amplicon of the supposedly *Pseudocercospora*-specific primers in the *Zasmidium syzygii* P124 genome assembly and in the *P. fijiensis* reference genome assembly (Cirad86; Arango et al., 2016). Both isolates possess the primer sequence used to distinguish *P. fijiensis* (Table S1) from *P. musae* and *P. eumusae* and produce a similar sized amplicon of 480 bp in *Z. syzygii* P124 and 478 bp in *P. fijiensis* Cirad86, which explains the positive result for *Z. syzygii* in our PCR assay (Figure 1). The amplicons share 88% sequence identity and *Zasmidium* or *Pseudocercospora* isolates can therefore be distinguished only upon amplicon sequencing. Thus, novel primer pairs need to be developed to enable easy and accurate diagnosis of fungal species present in foliar blights of banana.

## Conclusion

Here, we report the first chromosome-scale genome assembly of a *Zasmidium* species, this adds a high-quality genome sequence to the thus far limited genetic resources available for this genus. Our data show that *Z. syzygii* occurs on banana foliage in Bohol and can cause leaf necrosis, comparable to the foliar blight symptoms observed for *P. fijiensis*. The availability of the genome assembly will facilitate further research into the association of *Zasmidium, Pseudocercospora*, and possibly other fungal species related to foliar blights of banana. Moreover, it will serve as a valuable resource for developing novel molecular diagnostics, enabling the accurate identification and characterization of these fungal species.

## Data Availability Statement

All sequencing data are deposited in the NCBI database under accession number PRJNA931964. Scripts used for the genome assembly and analysis are available on github https://github.com/Anouk-vw/Zasmidium.

## Acknowledgments

We thank the Research Support Group (RSG) at KeyGene B.V. (Wageningen, the Netherlands) for performing the DNA isolation, library preparation and genome sequencing. We thank Xiaoqian Shi-Kunne (Wageningen University & Research, the Netherlands) for the data management and bioinformatics support.

