## Supplementary Material for "A chromosome-level genome assembly of *Zasmidium syzygii* isolated from banana leaves"

**Supplementary Table 1**. List of primers used in this study.

| Code | Name | Sequence (3’-5’) | Application |
|---|---|---|---|
| O1 | ACTF1 | TCCAACCGTGAGAAGATGAC | General for fungi |
| O2 | ACTR1 | GCAATGATCTTGACCTTCAT | General for fungi |
| O3 | Pf-actF | CTCATGAAGATCTTGGCTGAG | Specific for *P. fijiensis* |
| O4 | Pm-actF2 | ACGGCCAGGTCATCACT | Specific for *P. musicola* |
| O5 | Pm-actRb | GCGCATGGAAACATGA | Specific for *P. musicola* |
| O6 | Pe-actR | GAGTGCGCATGCGAG | Specific for *P. emusae* |

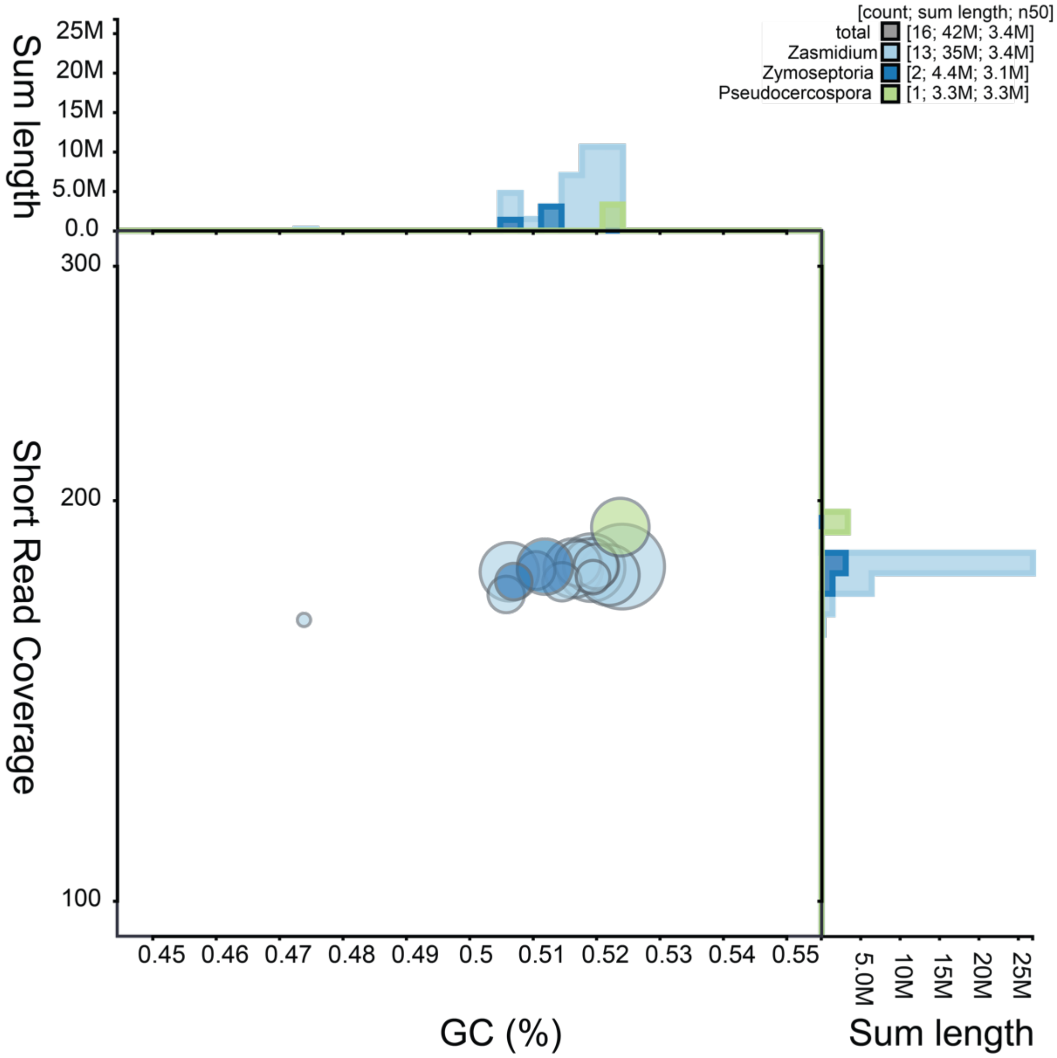


**Supplementary Figure 1.** Blob plot of base pairs found in the NCBI taxonomy database and the GC content. Fourteen contigs (35 Mb) have the best hits against species from the *Zasmidium* genus, the other 6.4 Mb of the genome have their best hit with other members of Mycosphaerella, Pseudocercospora (3.3 Mb) and Zymoseptoria (3.1 Mb). The GC content of the contigs is around 50% and the contigs are covered by 180 reads on average, suggesting that all contigs belong to a single species.

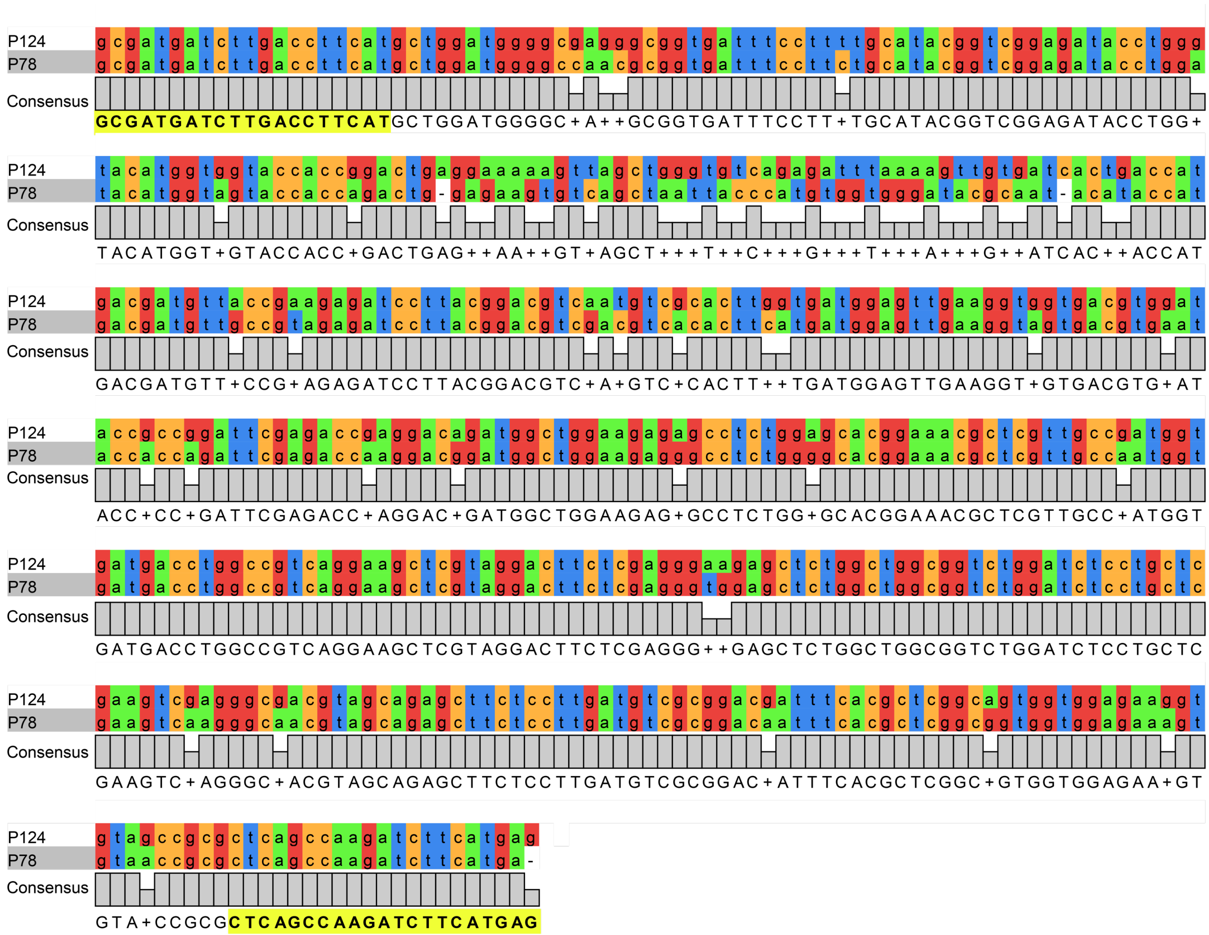


**Supplementary Figure 2.** Nucleotide sequence alignment of the PCR amplicon amplified using the *P. fijiensis*-specific primers. Primer sequence is highlighted in yellow.
